# Micro and Nanoplastics Interactions with Plant Species: Trends, Meta-Analysis, and Perspectives

**DOI:** 10.1101/2022.02.11.480069

**Authors:** Imran Azeem, Muhammad Adeel, Muhammad Arslan Ahmad, Noman Shakoor, Muhammad Zain, Naglaa Yousef, Zhao Yinghai, Kamran Azeem, Pingfan Zhou, Jason C. White, Xu Ming, Yukui Rui

## Abstract

The ubiquitous presence of nano plastics (NPx) and micro plastics (MPx) in the environment has been demonstrated, and as such, the exposure scenarios, mechanisms of plant response, and ultimate risk must be determined. However, the current literature reports ambiguous outcomes and provides limited mechanistic insight into critical governing processes. Here, we performed a meta-analysis of the most recent literature investigating the effect of MPx/NPx on plant species under laboratory and field conditions so as to evaluate the current state of knowledge. Toxic effects of MPx/NPx exposure in plants varies as a function of plant species and interestingly, generally non-significant responses are reported in staple crops. NPx (<100 nm) more negatively affected plant development parameters (n=341) (n is total number of observations), photosynthetic pigments (n=80), and biochemical indicators (n=91) than did MPx (>100 nm). Surprisingly, NPx exposure yielded negligible effects on germination rate (n=17), although root morphology (n=45) was negatively affected. Alternatively, MPx negatively affected on germination (n= 27) and generally non-significant affect with regard to root morphology (n=64). The effect of MPx/NPx on plant health decreases with increasing exposure time. No specific trends were evident for the production of biochemical enzymes as related to MPx/NPx concentration or size. Future work should include crop full life cycle studies to highlight the accumulation of MPx/NPx in edible tissues and also to investigate potential trophic transfer of MPx/NPx. Furthermore, we provide a framework for additional investigative work to address these and other knowledge gaps and to enable accurate assessment of the fate and risk of these materials to environmental and human health.

**Environmental significance:** Accumulation of plastic (MPx and NPx) particles is increasing in environmental compartments, and this might be threatened to agricultural plants.

**Graphical Abstract:** 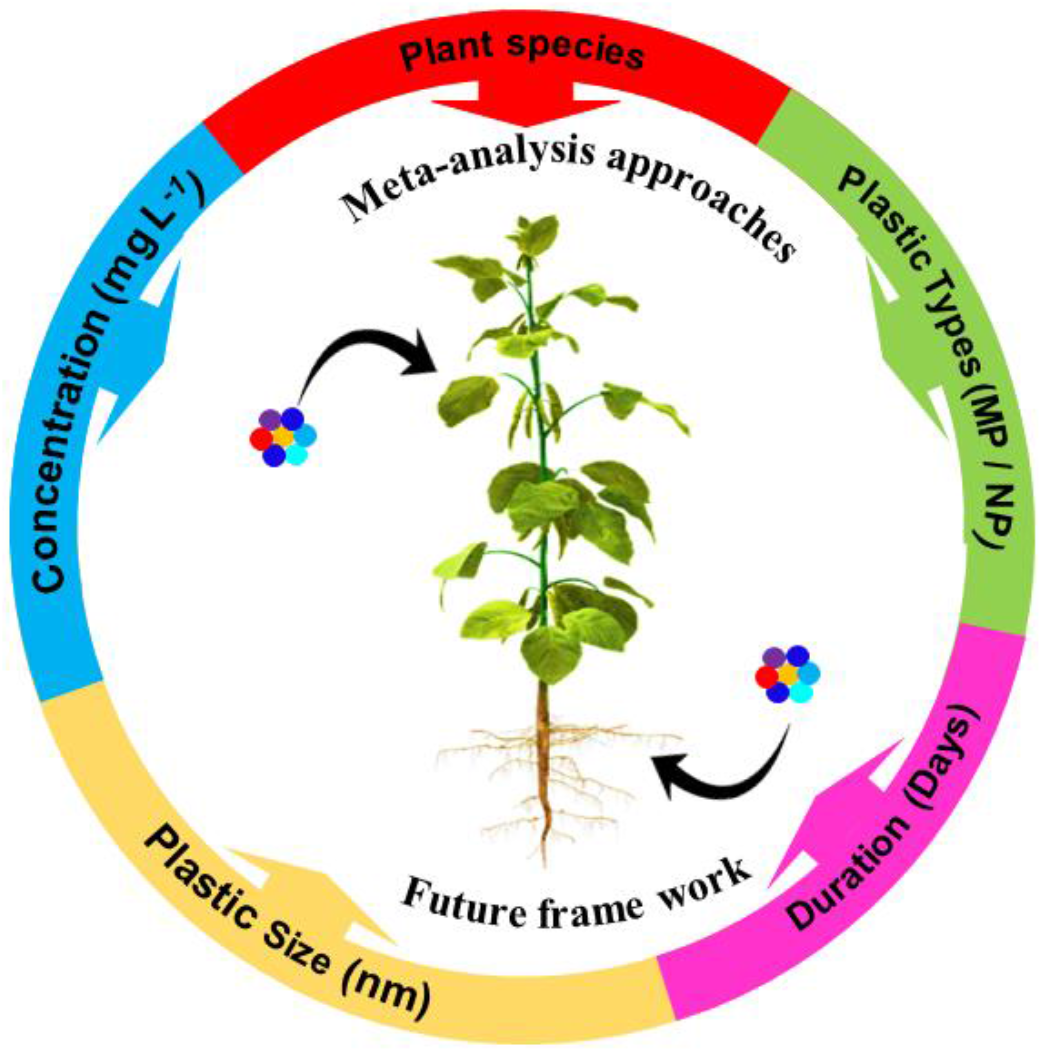

## 1. Introduction

Plastic contamination has become a widespread societal and ecological threat due to its global application, persistence, and adverse impact on biota.^1^ Global plastic production has reached 368 million tons, with production in China and Europe at 114.08 and 58.88 million tons, respectively; some predictions of global production to exceed 33 billion tons by 2050.^2–4^

Soil systems have become a major sink of plastics due to agricultural activities, including through amendments of sewage/sludge and organic fertilizers, as well as through the use of plastic films and mulches.^5–7^ Specifically, the application of municipal sludge in agriculture adds approximately 7.76 million tons of synthetic fibers or sedimented microplastic (MPx)/ nanoplastic (NPx) to soil systems each year.^8–10^ Furthermore, mulching use for crop production as part of weed control and irrigation strategies introduces approximately 4.4 million tons of plastics each year to soils.^11, 12^ Additionally, organic fertilizers from biowaste fermentation and composting have been reported to contain up to 895 particles kg^−1^ and have been identified as major source of plastics contamination of soil.^13^ In addition, landfill sites has been identified as one of the most significant entry pathways of plastic pollution into the environment.^14, 15^

A number of recent studies have investigated the effects of MPx/NPx exposure on crop species, with findings ranging from neutral effects to overt phytotoxicity.^16–20^ Despite the known exposure and accumulation scenarios in agricultural systems, understanding of the interactions of these materials with agricultural plants remains elusive. For example, Lian et al. ^21^ and Jiang et al ^22^ conducted detailed investigations of NPx exposure on wheat (*T. aestivum*) and broad bean (*V. faba*) and reported mixed findings. Lian et al. reported that polystyrene (PS) NPx had no significant effects on *T. aestivum* growth,^21^ whereas exposure of *V. faba* to PS-MP (50 and 100 mg L^−1^) significantly decreased (48 and 63%) plant biomass.^22^ Moreover, Sun et al. and Lian et al. reported deleterious effects of foliar application (1 mg L^−1^) of PS-NPx on *Lactuca sativa* (*L. sativa*) and *Zea mays* (*Z. mays*) in soil.^23, 24^ Recently, Azeem et al. and Yan et al. reported toxicity of MPx/NPx as measured by enzymatic activity and physiological parameters of the plants.^25, 26^ However, a mechanistic understanding of the role of material size, concentration and plant species-specific response to MPx/NPx on plants physiological, biochemical, and photosynthetic indices remains elusive. It is clear that investigations of the ecotoxicology of MPx/NPx are needed to gain a comprehensive assessment of their hazard (s) and ultimate risk in agroecosystems.

Given the known and expected discharge of plastics into the environment, we conducted a meta-analysis using the published literature on effects of MPx/NPx on agricultural plant species. We also focused on evaluating the significance of size and concentration of plastics on the physiological and biochemical response of different plant species. We also highlight research gaps, discuss methods, and offer recommendations and perspective for future research.

## 2. Methodology

### 2.1 Literature search

A total of 654 studies (prior to October 30, 2021) that investigated the interaction of MPx and NPx in the environment were collected from Web of Science, Scopus, Google Scholar, PubMed, and ResearchGate. A number of keywords were used for the search including “plastic particles”, “microplastic”, “nanoplastic”, “plants”, “field crops”, “physiological response” and “biological indicator” (Figure S1). We developed and executed a strict search strategy to collect the most relevant, novel, and reliable data sets (Figure 1). As noted above, the initial search returned 654 research articles and that group was narrowed to 25 articles by mandating the following criteria in our search strategy: (i) the study included the application of plastic (MPx or NPx) on agricultural plants (ii) the experiment was conducted under laboratory or field conditions (iii) a control or MPx/NPx-free treatment was included and (iv) the results were supported by appropriate statistical analyses. Only studies that met all four criteria were included for analysis.

**Figure 1.**
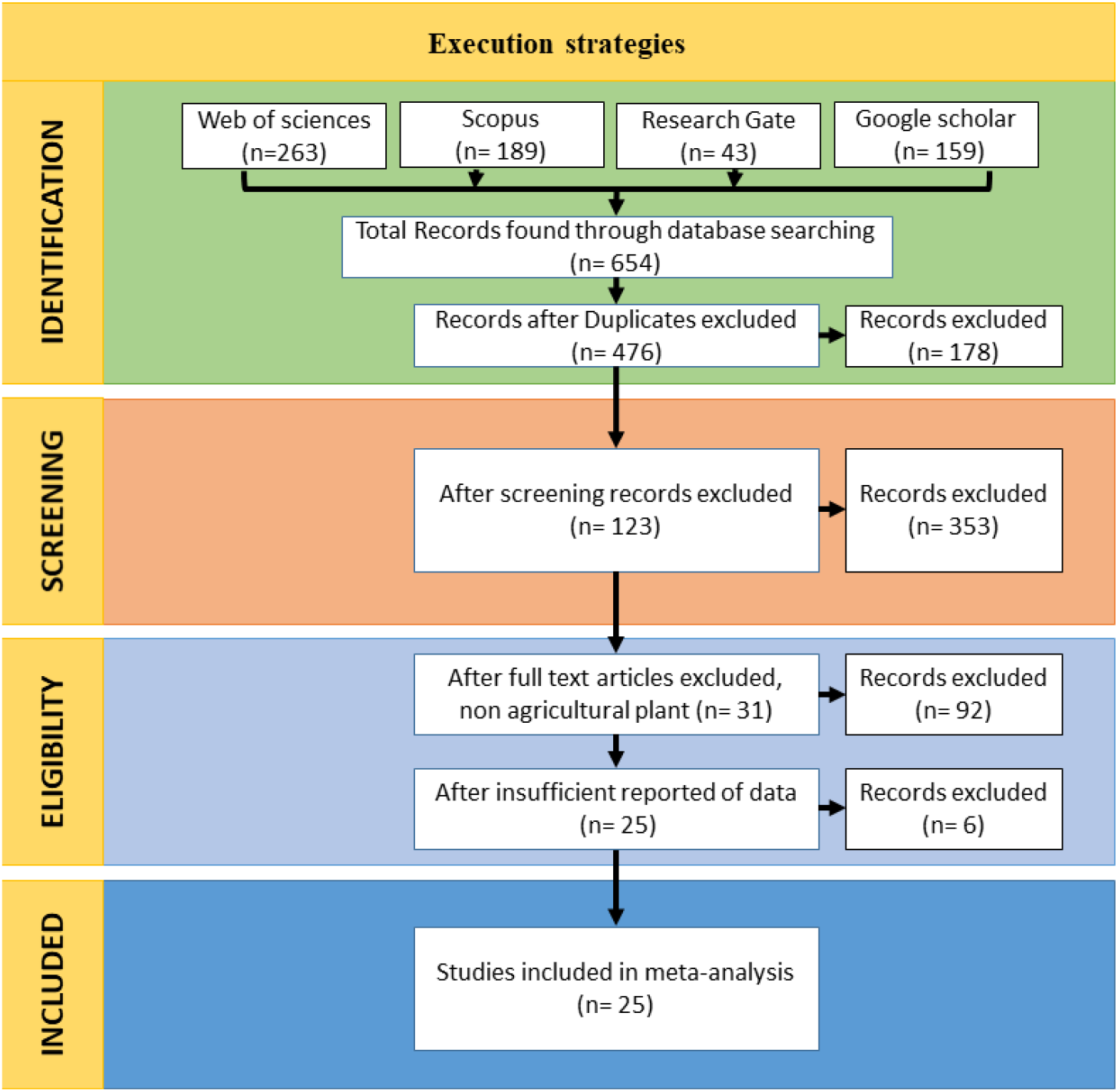
Work plan strategy for conducting the meta-analysis

### 2.2 Data extraction

Data was recovered from each study that met the above criteria, including plastic size and dose (Figure 2), plant species (Figure S3A), plastic type [polystyrene (PS), low-density polyethylene (LDPE), high-density polyethylene (HDPE), polylactic acid (PLA), fibers, polyethylene (PE), polyamide (PA), polyether-sulfone (PES), polyethylene terephthalate (PET), polypropylene (PP) and polymers coated fertilizer)] (Figure S3B). A total of 25 peer-reviewed research articles were included in the meta-analysis; the GetData Graph Digitizer software (version 2.26) was used to collect the data from the figures in published articles.

**Figure 2.**
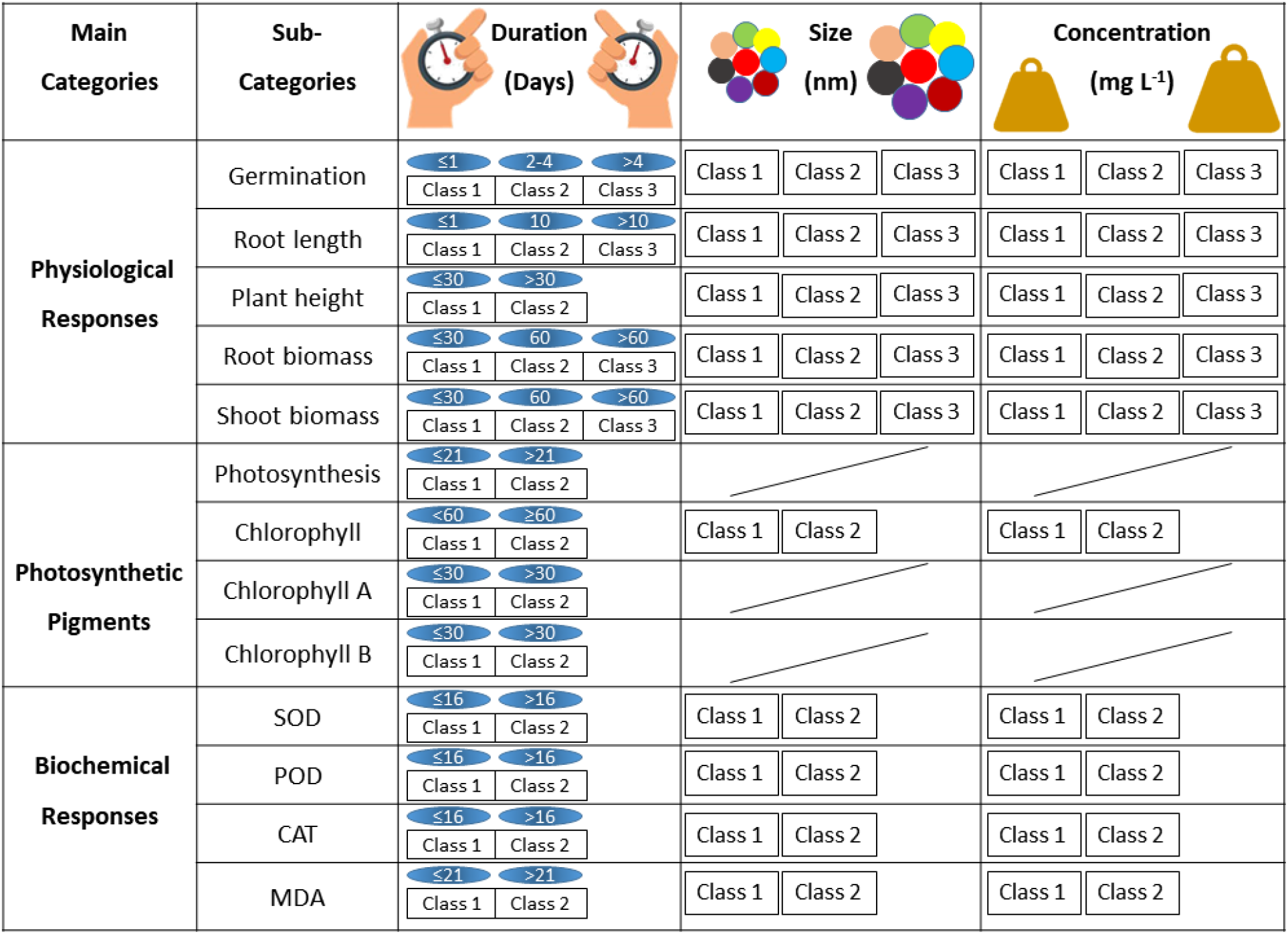
Meta-analysis classes with regard to MPx/NPx. The blue color shows the duration range; a line in a specific cell line indicates that there is no data. Duration is expressed in days, size is expressed in nm, and concentrations are in mg L^−1^.

### 2.3 Data endpoints

We classified all data endpoints (number of observations in the studies) into three main categories and thirteen subcategories depending upon their biological significance. Biological significance includes physiological response, photosynthetic pigments, and biochemical response (Figure 2). For example, if the authors of a particular study established different treatments of MPx or NPx and studied endpoints such as plant species, material size and type, and concentration, we included each endpoint separately for each treatment. The categories are shown in Figure 2.

Each of category or subcategory of an endpoint was expressed as a percentage, calculated by dividing affected endpoints by total number of endpoints in similar main or subcategories.

### 2.4 First-order meta-analysis

We used the natural log-transformed response ratio (lnRR) method as described by Hedges, et al. ^27^ and Gurevitch, et al. ^28^ to evaluate the effect of MPx and NPx size, type, and concentration against plant physiological response, photosynthetic pigment content, and biochemical parameters as shown in Figure 2. The following equation was used for estimating the effect size.

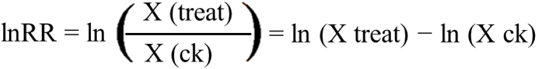

Where X (treat) was denoted for the mean of treatment, and X (CK) was denoted the mean of the control. If a study reported only standard error (SE), we calculated Standard deviation (SD) using SD = SE √(n) (n represented number of replicates). The estimated size effect was also converted to percentage through following equation = (elnRR-1) × 100%.

### 2.5 Statistical analysis

All statistical analyses and data preprocessing were performed in Microsoft Excel 2016, and R environment using meta^29^, metaphor ^30^ packages, and ggplot.^31^ The assessed polled effect was formed by first-order meta-analysis with 95% confidence intervals (95% CI) and is presented as forest plots. A negative value indicates a negative impact on plant parameters, and positive values indicate a positive effect.

## 3. Results and Discussions

### 3.1 Trends in research findings of MPx/NPx interactions with plants

We identified 25 research articles published prior to November 2021 that experimentally exposed plant species to MPx/NPx and met the criteria identified above. The first study was published in 2018; this emerging area of research has received increasing attention since then and published 11 research articles in 2021 (Figure S2). This trend will most likely continue to expand given the growing application and occurrence of plastic pollution at a global scale.^32^ Geographically, China is the leading country investigating the phytotoxicity of MPx/NPx on plants. Additional countries such as the Netherlands, Germany, Italy, and the United Kingdom have published articles that highlight the significant interactions between plastic types and plant species. Based on our analysis of the literature, fourteen research groups have published on the toxicity of MPx/NPx on plant species in the last five years.

### 3.2 Impacts of plastic type (MPx/NPx) on plant response

Generally, the plant response against MPx/NPx is presented in supplementary file (Result and discussion S1). However, in terms of plastic type, plant response to MPx exposure was studied more frequently than NPx. As measured endpoints, 368 (67%) focused on MPx compared to 182 (33%) endpoints with NPx (Figure 3). The meta-analysis results revealed that physiological endpoints were affected by MPx/NPx type; specifically, shoot biomass and root length were reduced by approximately 25%, with other decreases being evident for germination 13%, root biomass 13% and plant height 6%. The negative effect on seed germination of *L. sativum* was directly related to plastic type; germination with MPx (4800 nm) was only 21% as compared to 56% germination upon NPx (50 nm) exposure.^20^ *L. perenne* exposed to MPx (fiber and PLA) at 1 g kg^−1^ have reduced germination by 9 and 8% respectively.^16^ In summary, the meta-analysis findings show that NPx have a greater negative effect on root and shoot biomass than does MPx.

**Figure 3.**
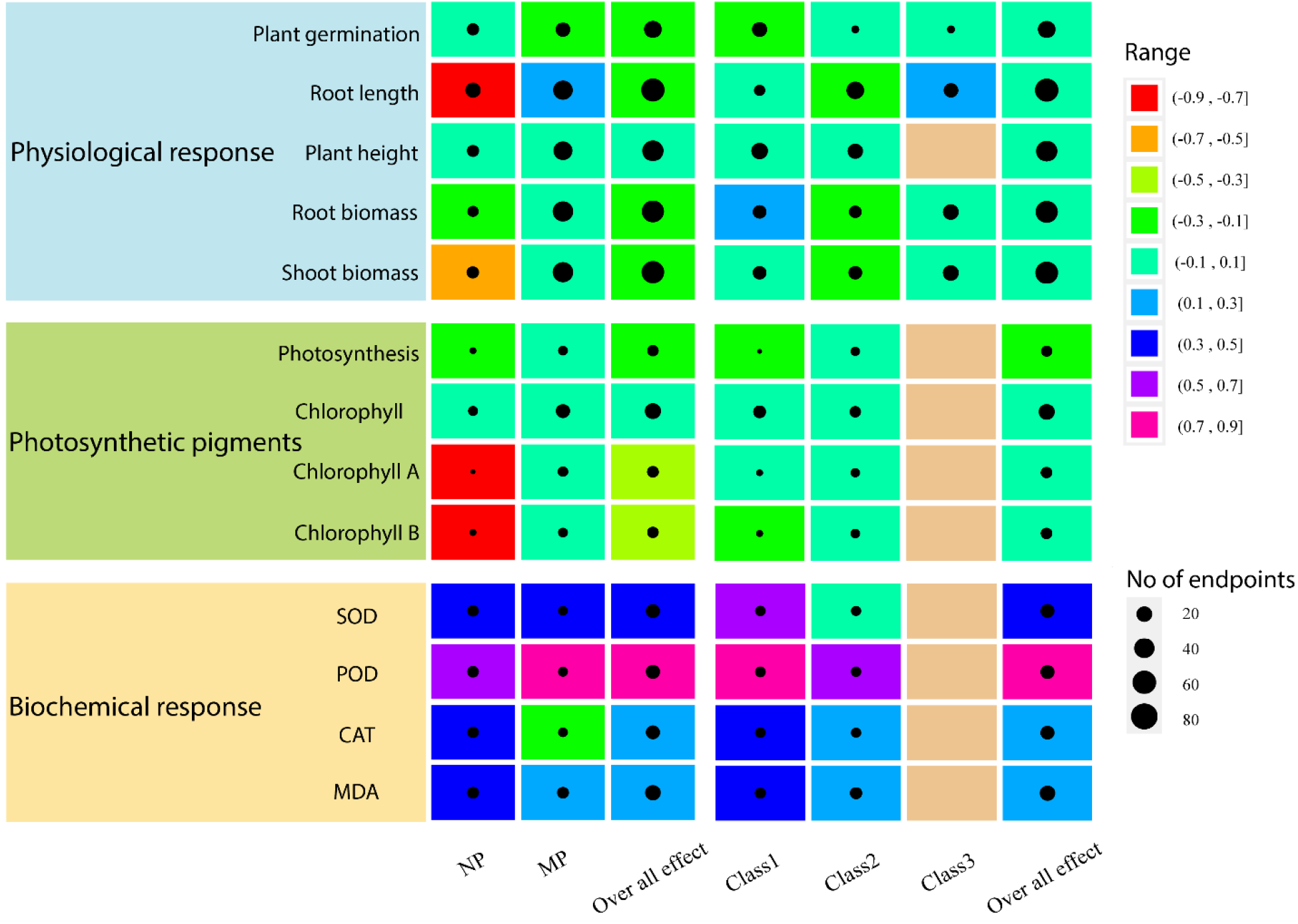
Eco-toxicological effects of plastic on plant biological functions according to their types (MPx, NPx, and overall effect) and duration (Class 1, 2, 3, and overall effect). The circle expresses the number of studies and color expresses the effect on plant biological function.

Exposure to MPx/NPx induced a negative response on photosynthetic pigments, with the most investigated reduction in chlorophyll b content (35%), chlorophyll a (32%) followed by photosynthetic rate (14%) and total chlorophyll content (9%) (Figure 3). The content of chlorophyll a and b in the leaves of *C. sativus* were reduced by 39% and 35% respectively upon PS-NPx, and while on an average at different doses MPx reduced the chlorophyll a by 9% and chlorophyll b by 11%.^33^ However, non-significant differences in relative chlorophyll content of *T. aestivum* were noted upon exposure to LDPE and biodegradable MPx at 10 g kg^−1^ ^18^; similar findings were reported with *Z. mays* and *L. sativa*.^32, 34, 35^ Importantly, the mechanisms driving the impacts, both negative and positive, on photosynthetic rate, total chlorophyll content and chlorophyll a and b with NPx/MPx exposure remain largely unknown.

Overall, MPx/NPx had notable effects on the biochemical activities of plants, with the most reported endpoints on POD with an increase of 104%, 47% increase in SOD, 15% in CAT and 32% in MDA activity. Interestingly, taking into account the plastic types, we have found that CAT activity was reduced by 20% with MPx exposure, while 64% increase was observed under NPx exposure (Figure 3). Jiang et al. observed that exposure of *Vica faba* to NPx increased the SOD (98%), POD (179%), CAT (57%) and MDA activity (15%), while PS-MPx application increased SOD (146%) and POD activity (324%), and CAT was decreased by 51% as compared to control, however non-significant changes were noted in MDA content.^36^ Similarly Li et al. reported that PS NPx (100 nm) increased SOD (23%) and CAT (22%) activities of *C. sativus*, while PS MPx (500 nm) enhanced SOD by 18% and decreased CAT by 63%.^33^ In short, NPx may have negative impacts than MPx on root length, root and shoot biomass; this appears to not be the case for germination and plant height. In addition, photosynthetic pigments were also reduced with NPx exposure than with MPx. Furthermore, exposure of MPx/NPx tended to increase antioxidant enzymatic activity, with the exception being CAT activity which was decreased upon MPx while increased under NPx exposure.

### 3.3 Effects of duration of exposure of MP/NP on plants

The duration of MPx/NPx exposure was highly variable among the twenty-five studies we analyzed, with a range of 8 hours to 120 days and an average of 22 days. With regard to physiological response, exposure for <1 day decreased germination by approximately 17%, while 2-4 and >4 days exposure has non-significant effect on germination. Root length (n=88) decreased under class 1 (<1 day exposure) and class 2 (10 days exposure) by 4% and 19%, respectively, while class 3 (>10 days exposure) significantly increased root length by 29%. Plant height (n=70) has two classes of exposure duration, i.e. 1 (<30 days) and 2 (30 days), and both showed negative response. The root and shoot biomass were enhanced with < 30 days exposure to plastic by 17% and 4%, respectively. However, 31-60 days exposure inhibited the root biomass (14%) and shoot biomass (18%), while >60 days exposure decreased the root and shoot biomass by 2% and 6%, respectively (Figure 3).

Exposure of <1 day to MPx/NPx clearly had a significant impact on germination; for example, NPx/MPx reduced the germination of *Lepidium sativum* with an exposure period of less than 24 hours, but after 24 hours, there was no effect.^20^ Similarly, non-significant effects were found in the *A. cepa* upon 3-days exposure to PS-NPx.^37^

Interestingly, exposure to NPx resulted in a significant reduction (16%) in root length of *L. sativum* after 24 hours, while with increasing time (after 48 hours) non-significant differences were observed.^20^ Conversely, Sun et al. reported that *Arabidopsis thaliana* root length was significantly decreased with PS (SO_3_H and NH_2_) exposure at 10, 50, and 100 mg L^−1^ after 10 days exposure.^38^ Alternatively in Class 3 (>10 days exposure), the root length of *A. fistulosum* was increased by MPx (PA, HDPE, PES, PET, PP and PS) after 45 days of exposure.^39^

The effect of MPx/NPx on root and shoot biomass of *T. aestivum* were evaluated after 30 days of exposure *and T. aestivum* had remarkably increased root and shoot biomass with PS NPx (0.01 and 0.1 mg L^−1^) after a 21-day exposure.^40^ Conversely, the exposure of MPx for 60 days inhibited the root (40%) and shoot biomass (55 %) in *T. aestivum*.^18^ While, more than 60 days exposure of bio degradable-MPx to *P. vulgaris* notably inhibited (21 and 18%) root and shoot biomass respectively at 15 g kg^−1^.^41^ From the meta-analysis results, it is clear that both plant species and duration are important parameters affecting the extent of MPx/NPx toxicity.

With regard to photosynthetic endpoints, different results (beneficial, harmful and neutral effects) were observed as a function of different exposure times (Figure 3). In general, the photosynthetic activity (n=15) was decreased by 14%, while in case of class based description, <21 days exposure decreased activity by 17%, and 6% reduction was observed upon >21 days exposure. Chlorophyll a (n=16), <30 days exposure led to increased (9%) content but >30 days exposure resulted in a 1% reduction. For chlorophyll b (n=16), <30 days exposure decreased content (14%) as compared to >30 days exposure (Figure 3).

Application of PS NPx decreased the photosynthetic output of *L. sativa* after 21 days of exposure.^24^ Zeb et al. treated *L. sativum* plants with PS microfiber for 58 days and found an 11% reduction in chlorophyll contents at 20 g kg^−1^ concentration.^35^ Exposure to biodegradable-MPx (15 g kg^−1^) for 103 days decreased chlorophyll content (13 %) in *P. vulgaris*.^41^ *Lolium perenne* at 30 days of exposure to MPx (fiber, HDPE and PLA) increased the chlorophyll a content by 3, 9 and 17% but decreased chlorophyll b levels by 6, 15 and 21%, respectively.^16^ Overall, in class 1 (<21 days for photosynthesis and <60 days for chlorophyll), photosynthesis and chlorophyll b were negatively affected as compared to the class 2 exposure (>21 days for photosynthesis and >60 days for chlorophyll). Decreased chlorophyll content may be attributed to metabolic dysfunction or to the accumulation of ROS in chloroplasts, ^42^ which are important sites for ROS generation under stress condition.^43^ However, more detailed mechanistic studies are needed to better understand the impact of different types of MPx and NPx under different exposure times on the photosynthetic activity and pigments of different plant species.

In terms of biochemical response, the overall impact of exposure duration on biochemical activities as determined by SOD (n=25), POD (n=25), CAT (n=25) and MDA (n=32) were significantly increased by 35%, 107%, 29%, and 40%, respectively (Figure 3). Jiang et al. reported that *V. faba* under 2 days exposure of PS MPx (5 μm) had significantly increased SOD and POD activity, while CAT activity was decreased with MPx and increased with NPx (100 nm). While, MDA activity was unaffected by MPx, whereas NPx at 100 mg L^−1^ enhanced MDA activity.^36^ The application of PS-NPx for 2 and 7 days significantly increased the SOD, POD, CAT and MDA activity by 201, 110,1 92 and 176%, respectively in *Z. mays*.^44^ Similar results on SOD, POD and CAT activity (75, 97 and 55% respectively) were found in *O. sativa* after 16 day of PS-NPx exposure.^45^ In *C. sativus*, 65 day exposure of NPx (100 nm) increased SOD, POD and CAT activity (27, 333 and 22% respectively); interestingly, MPx (500 and 700 nm) decreased SOD (19 and 74%) and CAT (63 and 59%) activities, but increased (50 and 238%) POD at the 300 and 700 nm size.^33^ We observed that class 1 (<16 days exposure) generally increased the enzymatic activity of plants as compared to class 2 (>16 days exposure). The mechanisms for this difference are unknown but these materials might be associated with the growth media, root exudates or other plant metabolites that reduced the overall toxic effect of MPx/NPx; further investigations are needed here. In addition, increasing the MPx/NPx exposure time generally induces a decrease in antioxidant enzymatic activity. However, more mechanistic full life cycle studies are needed to understand underlying mechanisms mediating the interaction between MPx/NPx exposure duration and plant biochemical response.

### 3.4 Size effects of MPx/NPx on plant

Depending on endpoints, the smaller size plastic particles (i.e., <5 μm) are generally assumed to be more toxic as compared to larger size materials.^25, 46^ A negative response on plant germination was resulted from exposure to MPx/NPx and this response varied with size. With MPx/NPx sizes of <100 nm, 100-400 and >400 nm, germination decreased by 7%, 19% and 27%, respectively. Small plastic size particles (<100 nm) significantly decreased (9%) the root length, while large size plastic particles (>500 nm) have shown a positive effect (22%) on root elongation (Figure 4). For plant height, MPx/NPx sizes of <100, 100-400, and >400 nm showed non-significant effects. Bosker et al. reported that the germination *of L. sativum* was reduced (31%) at 50 nm plastic size, but the greatest reduction (42 and 54%) on germination was at 500 and 4800 nm plastic size respectively. Furthermore, Boots et al. found that large size plastic (>400 nm) significantly reduced germination and height by 7 and 19%, respectively in *Lolium perenne*.^16^

**Figure 4.**
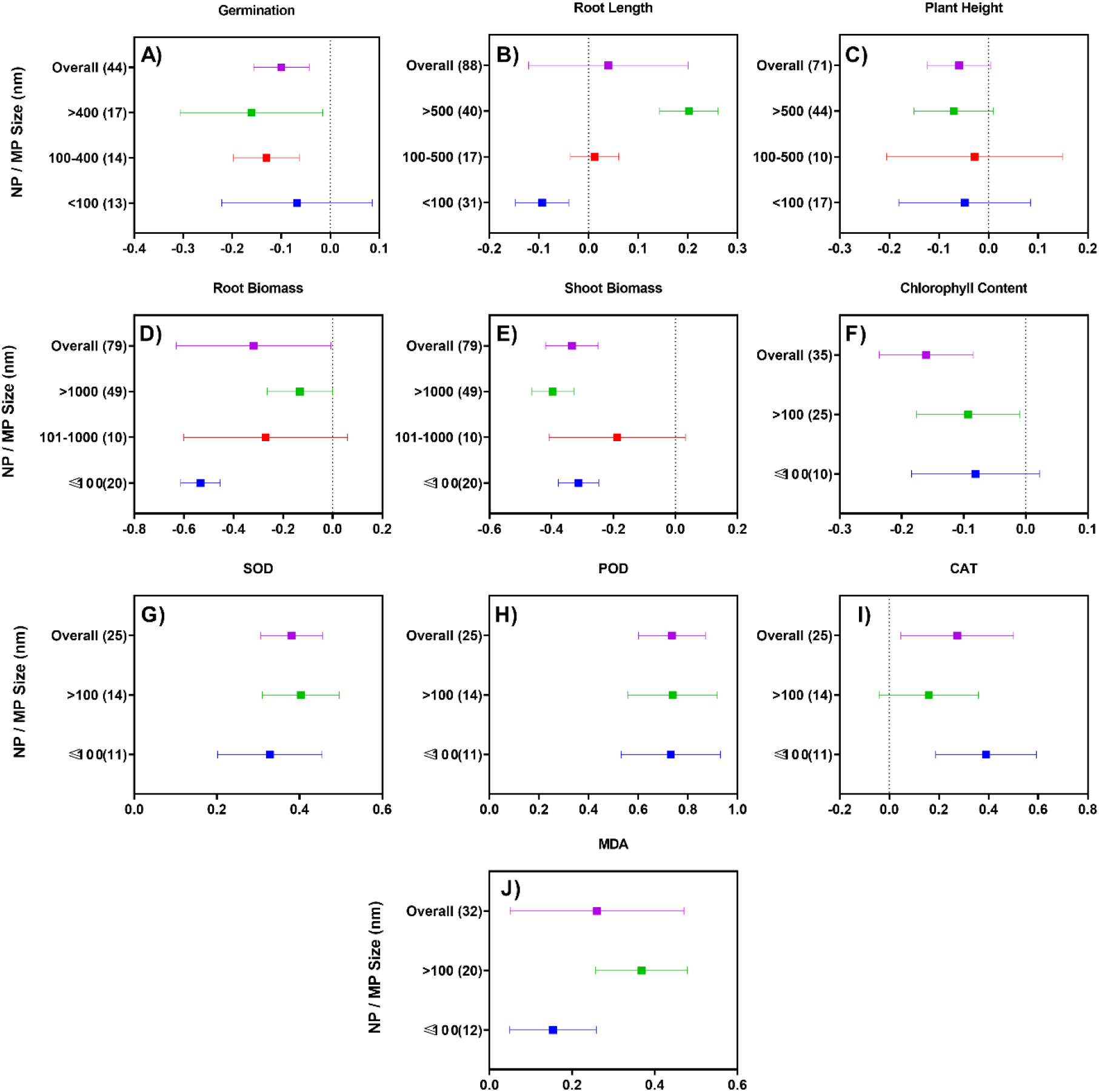
Mean effect of plastics size on germination (A), root length (B), plant height (C), root biomass (D), shoot biomass (E), total chlorophyll content (F), SOD (G), POD (H), CAT (I) and MDA (J). Sample size is noted on the left side bars, and the corresponding number of research studies is shown in parentheses.

MPx/NPx size exerted generally negative impacts on root biomass. For example, root biomass was significantly reduced by 41% with ≤100 nm size particles, followed by 24% in 101-1000 nm and 12% with MPx of >1000 nm. Shoot biomass showed a similar negative response under various sizes of MPx/NPx particles. Significant negative reductions (27%) in shoot mass were observed at <100 nm, followed by 17% at 101-1000 nm, and 34% at >1000 nm. Sun et al. reported that *Arabidopsis* biomass was significantly decreased by 41.7 and 51.5% upon exposure to 55 nm and 71 nm of PS plastic, respectively.^38^ Wang et al. reported that exposure of PLA plastic (100 μm size) reduced *Z. mays* root and shoot biomass by 69% and 53% respectively. *C. sativus* shoot biomass was reduced (48 and 25%) by exposure to 300 nm, and 700 nm PS plastic respectively. However, root biomass was reduced by 50% at 100 nm and 300 nm had no effect.^33^ In *P. vulgaris*, biodegradable plastic (250-500 μm size) reduced the shoot biomass by 25% but root biomass was increased (11%).^41^ PS plastic of 93 nm size reduced *L. sativa* total dry biomass by 23%.^24^ Our meta-analysis highlights that root and shoot biomass were directly affected by MPx/NPx size but this varies significantly with plant species. Additional studies are required to understand the long-term effects of various sizes of MPx/NPx on plant species, as well as the underlying mechanisms that drive those responses as a function of material size.

In our meta-analysis findings, total chlorophyll content showed generally non-significant effects from exposure to MPx/NPx, with reductions of 8% at ≤100 and 9% at >100 nm. PS small size plastic particles (0.071 μm) reduced the chlorophyll content of *A. thaliana* by 21%.^38^ While PS large size plastic particles (255 μm) decreased the chlorophyll content of *L. sativa* by 10.8%.^35^ By contrast, *T. aestivum* treated with PS small size plastic particles (100 nm) had a 22.7% increase in the total chlorophyll content.^47^ However, Qi et al. reported that *T. aestivum* exposed to LDPE and biodegradable large size plastic particle (5 μm) had no impact on chlorophyll content.^18^

Plant SOD activity was markedly affected by NPx/MPx as a function of size, as significant increases were observed at ≤100 nm (36%) and at >100 nm (50%) size. Plant POD activity was also increased with MPx/NPx size, including 108% at ≤100 and 109% at >100 nm particle size. Additionally, CAT activity was increased by 48% upon exposure of ≤100 nm and 17% increase was observed with particles of >100 nm. A similar trend was observed with MDA activity specifically, <100 nm increased values by 17% and under >100 nm the increase was 44%.

The SOD, POD and CAT enzymatic activity in *O. sativa* was increased upon exposure of PS small size plastic particles (20 nm).^45^ Similar results were found with *Vicia faba*, where SOD, POD, CAT, and MDA activity were all increased upon exposure to 100 nm (98%, 179%, 57% and 15% respectively) and at 5000 nm plastic size SOD and POD were increased by 146% and 324% respectively, while MDA had no effect; however, 51% reduction was observed in CAT activity.^36^ Zeb et al. reported that exposure of *L. sativa* to 255 μm size plastic particles increased SOD, POD and CAT activities by 19, 18, and 12% respectively and decreased (9%) the MDA activity.^35^

### 3.5 Effects of MPx/NPx dose

The MPx/NPx concentration (n=44) had a significant impact on plant germination. Our meta-analysis findings showed that MPx/NPx inhibited germination at ≤1 mg L^−1^ by 14%, 2-10 mg L^−1^ (13%) and >10 mg L^−1^ (9%) (Figure 5A). In case of root length, low concentrations (≤1 mg L^−1^) (n=33), exhibited a highly negative response (39% reduction), at medium concentration (n=25) values were less effected (15% reduction), and interestingly, at high concentrations (n=30) a positive response was evident (239% increased) (Figure 5B). For example, De Souse Machado et al. reported that *A. fistulosum* root length was increased by 25% on exposure to NPx/MPx at 20 g Kg^−1^.^39^ Zeb et al. observed similar findings with *L. sativa* by treating with PS-MPx at 10 and 20 g Kg^−1^and reported increase in 32% and 31% respectively. ^35^ Conversely, the root length of *O. sativa* was inhibited by 13% and 22% with PS-NPx at 50 and 100 mg L^−1^.^45^

**Figure 5.**
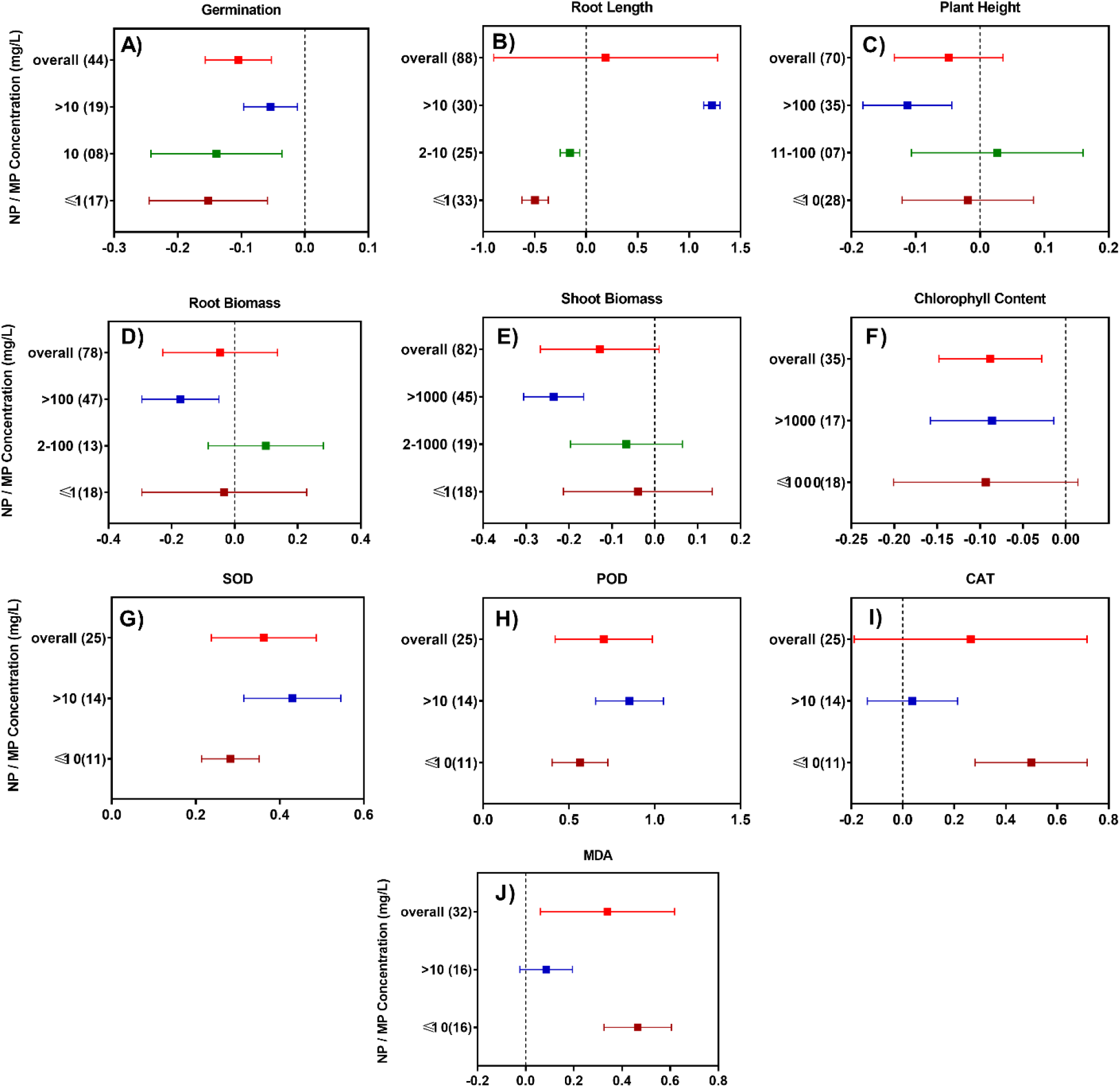
Mean effect of plastic concentration on germination (A), root length (B), plant height (C), root biomass (D), shoot biomass (E), total chlorophyll contents (F), SOD (G), POD (H), CAT (I) and MDA (J). Concentrations are noted lift side bars, and the corresponding number of research studies is shown within the parentheses.

Exposure to MPx/NPx showed negative effects on plant height (n=70) (Figure 5C) and root biomass (n=78) as a function of concentration (Figure 5D). Our meta-analysis shows that plant height and root biomass were decreased by 2% and 3% at <10 mg L^−1^, and increased by 3% and 10% at 11-100 mg L^−1^, while 11% and 16% reduction was found at >100 mg L^−1^ respectively. Exposure to MPx/NPx had dose dependent negative effects on shoot biomass (n=82) (Figure 5E). Shoot biomass reduced by 4% at ≤1 mg L^−1^ concentration, and 6% at 2-1000 mg L^−1^, although at higher concentrations (>1000 mg L^−1^), increases (21%) in shoot biomass were evident. Lian et al. reported that *L. sativa* plant height showed 14% decreases at 0.1 mg L^−1^, with even greater reductions (23%) at 1 mg L^−1^ of PS-NPx, similarly total shoot biomass was decreased by 27% and 24% at both concentrations respectively.^24^ Generally, our analysis shows that low (class 1) and medium (Class 2) MPx and NPx concentrations under each endpoint have non-significant effects on plants height, root and shoot biomass, whereas high concentration tend to be of more significance with regard to physiological parameters.

By taking average at all concentration and classes (n=35), the total chlorophyll content was decreased by 8% (Figure 5F). PS-NPx reduced the chlorophyll content at 1000 mg L^−1^ but at 300 mg L^−1^ contents were equivalent to the control.^38^ PS-MPx decreased (10%) the chlorophyll content in *L. sativa* at both 1000 mg L^−1^ and 2000 mg L^−1^.^35^ Alternatively, *T. aestivum* total chlorophyll content was unaffected by exposure to PS-NPx exposure at 10 mg L^−1^.^47^

With increasing dose of MPx and NPx, the activities of SOD (Figure 5G) and POD (Figure 5H) enzymes were increased. At low concentration (≤10 mg L^−1^), the increases in SOD and POD activities were 35 and 76%, respectively; at higher concentration (>10 mg L^−1^), the increases were greater (54 and 135%), respectively. The activities of CAT (Figure 5I) and MDA (Figure 5J) were also increased upon exposure to MPx/NPx. At low concentration (≤10 mg L^−1^), the levels of CAT and MDA were increased by 65% and 59%, respectively, as compared to control, but at higher concentrations (>10 mg L^−1^), only 4% and 9% increase were noted.

Zhou et al. reported that 50 mg L^−1^ and 100 mg L^−1^ of PS-NPx notably increased POD activity (43% and 97% respectively) and SOD (60 and 75% respectively) in *O. sativa* plants.^45^ Jiang et al. showed that exposure of MPx at 10, 50 and 100 mg L^−1^ increased SOD (68, 141 and 229% respectively) and POD (297, 333 and 342% respectively) in *V. faba*, while the corresponding values were (30, 187 and 77% respectively) and (173, 369 and −7% respectively) upon exposure to NPx at 10, 50 and 100 mg L^−1^ respectively. Furthermore, CAT activity was reduced (13, 60 and 80%) by MPx at 10, 50 and 100 mg L^−1^ but was increased (75, 85 and 11%) by NPx at 10 mg L^−1^, 50 and 100 mg L^−1^.^36^ In general, the meta-analysis results highlighted that physiological indicators were reduced with concentration of MPx/NPx, excluding root length which was increased with concentration; and impacts on chlorophyll contents were generally minimal. Antioxidant enzymes were increased with MPx/NPx concentration, with the exceptions being CAT and MDA activities which were negatively correlated with concentration.

### Research gaps and perspective

Our current meta-analysis results showed that MPx/NPx affected a broad range of endpoints in over two dozen studies and suggest toxicity, metabolic dysfunction of MPx/NPx exposure to agricultural plant species. Importantly, the literature is both limited and highly variable in terms of design, making the existing data set an inadequate basis to fully assess the risk of MPx/NPx exposure. Based on the above observations and current analysis, the following priorities are recommended for future investigations:

1. A more thorough understanding of the mechanisms of MPx/NPx phytotoxicity is needed through the use of transcriptomic, proteomic and metabolomic endpoints. Importantly, these parameters should be measured over time so as to understand the dynamics of plant response and resilience.
2. There is dire need to highlight the importance of MPx/NPx structure, composition, and size. This could likely be achieved through the establishment of a reference library of materials that would be available for use by the research community working in this area.
3. The dose-dependent response is needed to achieve, including both short- and long-term toxicity, fate and disposition of these materials in complex systems at a dose range that spans orders of magnitude. In addition, differential exposure scenarios must be systematically evaluated and ranked in terms of risk, including exposure via soil, hydroponic and foliar routes.
4. The interaction of MPx/NPx with other environmental contaminants under different growth conditions needs to be systematically investigated. Although our meta-analysis shows that MPx/NPx exposure alone is of concern, the ability of these materials to transport additional xenobiotics in biological systems remains a significant area of concern for exposure and risk.
5. The ability of MPx/NPx to move through trophic levels in the food web and to induce transgenerational impacts must be evaluated. This includes an assessment of human food chain contamination and exposure.
6. In the context of MPx/NPx waste management, it is necessary to develop sustainable remediation technologies. The phytoremediation of MPx/NPx by using plants selected through bioprospecting in soil ecosystems can also be a sustainable and eco-friendly option, particularly for scenarios where the plastics are associated with co-contaminants. The application of organic amendments or biochar to remediate MPx and NPx pollution should be investigated.
7. Root exudates and mucilage are major barriers to pollutants accumulation by plants. Sun et al.^38^ reported that PS-NH2 enthused the root secretes high level of exudates which disturbed the stability of PS and decreased plastic uptake in *Arabidopsis thaliana*. Detailed investigations should be focused on screening the plant species that may have the ability to degrade plastics through root exudation or specific rhizosphere microbial communities.

It is also important to note that much of the existing literature includes studies that have doses orders of magnitude beyond that expected in the environment. We need to focus on generating hard data that may increase our understanding of exposure and risk under environmental relevance conditions. MPx/NPx in particular presents more complication, as the mitigation of their adverse effects are probably not targeting the MPx/NPx themselves, but rather on secondary aspects like plastic design, use and waste-management strategies. Solid waste management, green chemistry, and efforts to create a truly circular economy are independent fields and if integrated together with regulatory action, can be helpful to reduce the burden of plastics in the environment.

4.

