## Supplementary Data for "Micro and Nanoplastics Interactions with Plant Species: Trends, Meta-Analysis, and Perspectives"

### Supplementary information

### Material and Methods

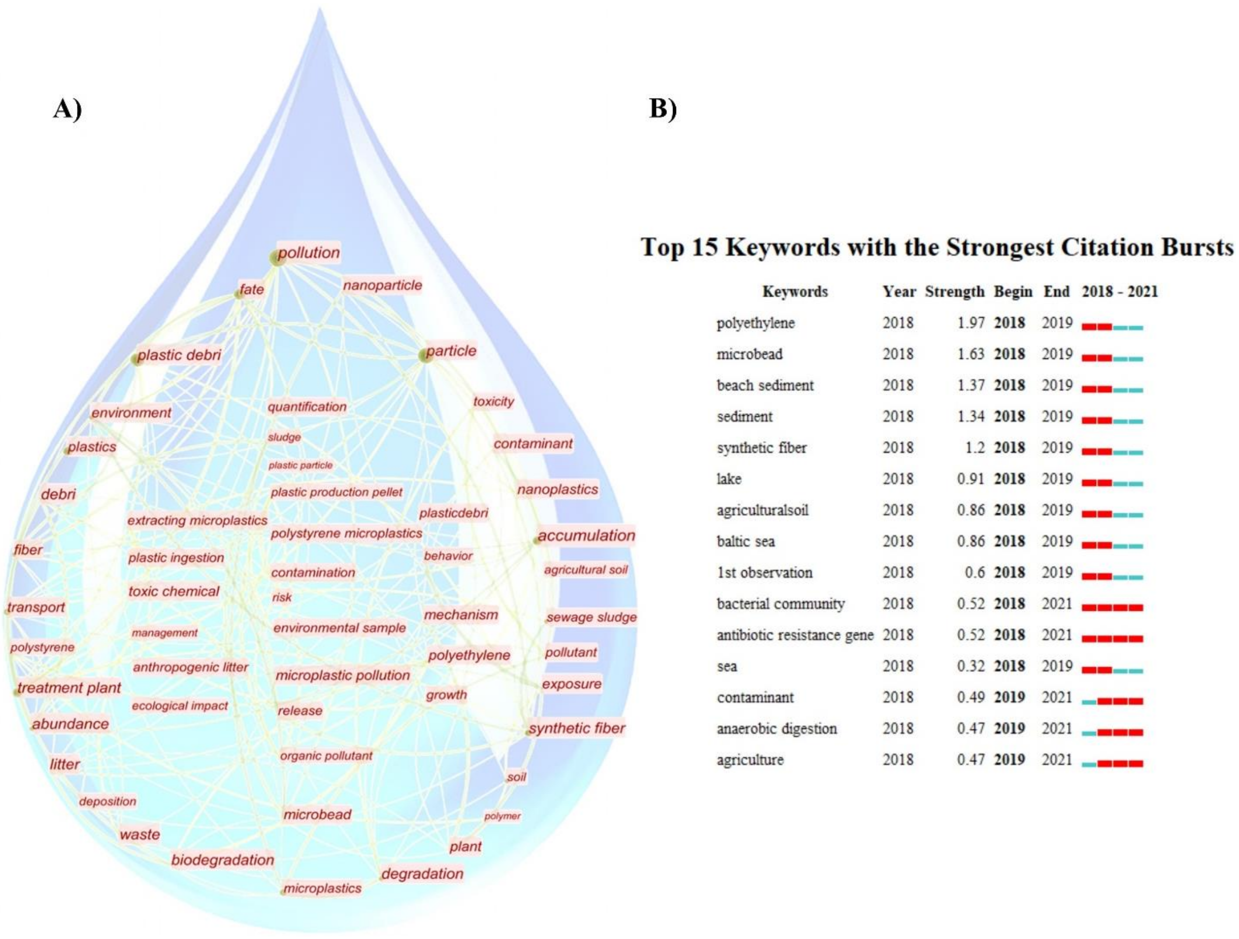


**Figure S1**. (A) The chart of high frequency keywords with time and keywords clustering from January 2018 to December 2021. The size of node indicates the frequency of keyword occurrence in the studied documents. (B) The top 15 keywords with the highest citation activity.

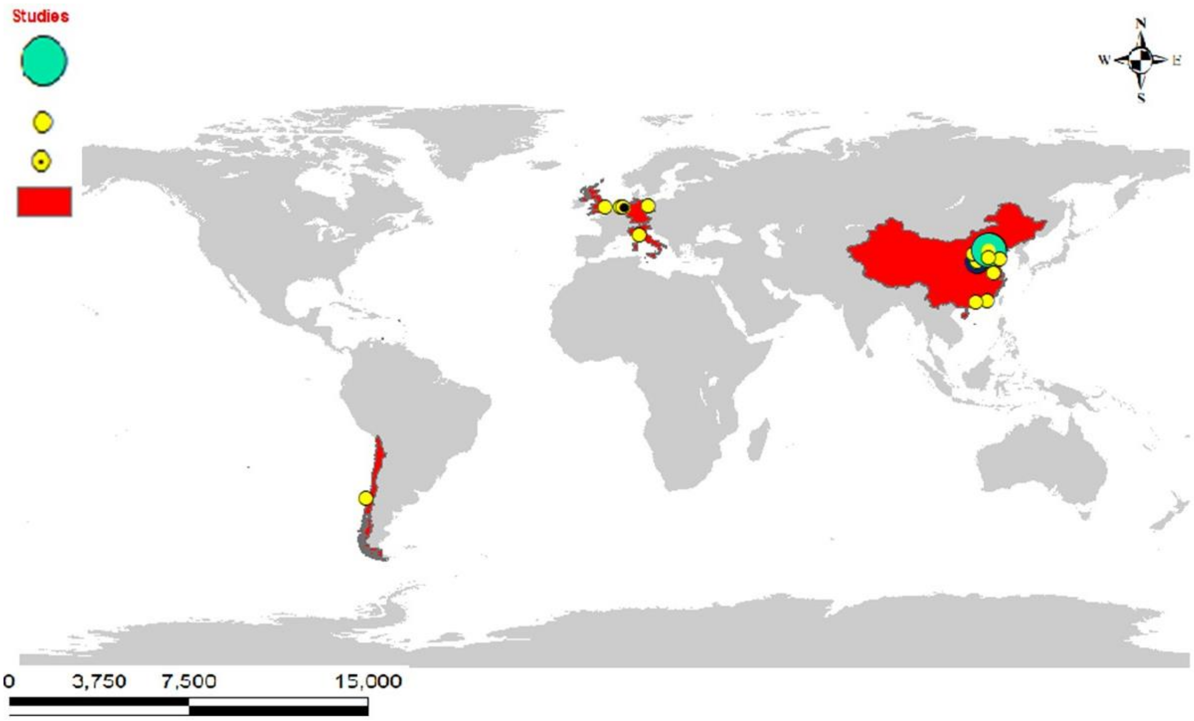


**Figure S2.** Published research articles which highlight the interaction of MPx/NPx with plants, red color indicates the country where the work has been done and remaining color shows the numbers of studies.

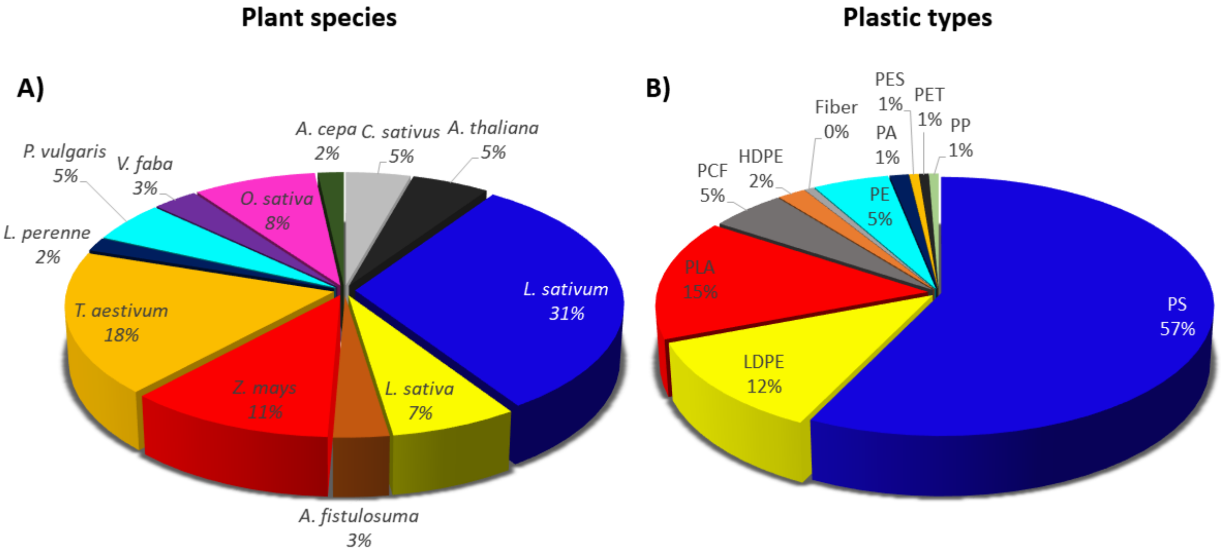


**Figure S3.** Published literature regarding the effects of MPx/NPx on plant species. (A) Plant species investigated (B) MPx/NPx types evaluated.

### Results and discussions

#### 1. Plant response against MPx/NPx

Agricultural crops are a primary component of human and animal diets.[1] Therefore, it is critical to investigate and understand the impact of different kinds of plastic on crops for both food safety and security. In this study, we analyzed the beneficial/harmful effects of MPx/NPx on individual agricultural plant species, with a focus on the endpoints shown in Figure S4. The most frequently measured endpoint category was a physiological response (n=341 endpoints), followed by biochemical (n=91) and photosynthetic response (n=80) (Figure S4).

Physiological response: In our analysis on the base of InRR, the endpoints of plant germination (n=44), plant height (n=70), and shoot biomass (n=70) were non-significantly reduced (6%, 8%, and 9%) upon exposure to MPx/NPx, whereas root length (n=87) and biomass (n=70) each were non-significantly increased by 3% (Figure S4).

Photosynthetic pigments: The endpoint of photosynthetic pigments shows different responses upon exposure to MPx/NPx. Photosynthetic rate (n=13), total chlorophyll content (n=35) and chlorophyll b (n=16) were reduced by 10%, 8% and 6%, respectively, whereas chlorophyll a (n=16) was unaffected by exposure (Figure S4). Importantly, chlorophyll a and b were only measured in four papers focused on exposure to MPx/NPx (n=45).[2-5]

Biochemical response: The biochemical response of plant species was consistently increased with MPx/NPx exposure; specifically, peroxidase (POD) (n=21) was increased 74%, superoxide dismutase (SOD) by (n=17) 56%, catalase (CAT) (n=21) by 27%, and malonaldehyde (MDA) (n=32) by 29% (Figure S4).

As noted above, there are published review articles describing the effects of MPx and NPx on plant species,[6] but the literature has not been systematically analyzed by plant species, plastic type, exposure duration, size, and concentration of MPx/NPx. Notably, in current meta-analysis, crop species are both positively or negatively impacted by exposure of MPx or NPx, depending one conditions; a detailed discussion of this analysis follows below.

Physiological responses such as germination, root length, plant height, shoot, and root biomass are widely used indicators of plant growth under stress exposure.[7] The current analysis confirms that exposure to various MPx/NPx can lead to a negative impact on plant species, with some exceptions such as *T. aestivum*, *Z. mays*, and *Oryza sativa* (*O. sativa*). For example, *Arabidopsis thaliana* (*A. thaliana*) and *V. faba* show more sensitivity to root length with values of

-40% and -21%, respectively, upon exposure of MPx/NPx (Figure S4). Furthermore, *Lactuca sativa*, *Cucumis sativus* (*C. sativus*), *P. valgaris* and *L. perenne* all demonstrated overall negative effects in response to plastic exposure. Interestingly, the root length *L. sativus*, *P. valgaris*, and *L. sativum* did exhibit a positive response[8], although the remaining species had a negative response to MPx/NPx.[9, 10] For example, *T. aestivum* germination was unaffected by PS-NPx, while a 190-220%, 70-87% and 52-98% increase was observed in root elongation, root, and shoot biomass respectively.[11] Conversely, Qi et al. reported that plant biomass was decreased by 38.9% upon exposure to MPx LDPE and biodegradable plastic at 10 g $kg^{-1}$.[12] Notably, findings in these studies seems to differ significantly as a function types and size of MPx/NPx. So, more studies are needed that highlight underlying mechanism of positive or negative effect of MPx/NPx on root length.

Chlorophyll a and b content are fundamental indicators of photosynthetic activity, with reductions in content or functionality often used as an indicator of plant stress and health.[13] In general, the total chlorophyll content and the content of chlorophyll b showed a negative response to MPx/NPx exposure, whereas chlorophyll a was generally non-significantly impacted (Figure S4). For example, Lian et al. reported that pigment content and photosynthetic response of *L. sativa* was inhibited upon exposure to PS-NPx at 1 mg $L^{-1}$.[2] Furthermore, PS-NPx disturbed chloroplast structure and photosynthetic activity, subsequently decreasing chlorophyll content.[2] Wang et al. reported that PLA-MPx significantly reduced the chlorophyll content at a higher dose (10, 100 g $kg^{-1}$).[14] However, there were no overt alterations in the chlorophyll a content of *L. perenne* upon 30 days exposure to various MPx (HDPE and PLA) at 1 g $kg^{-1}$ in soil.[4]

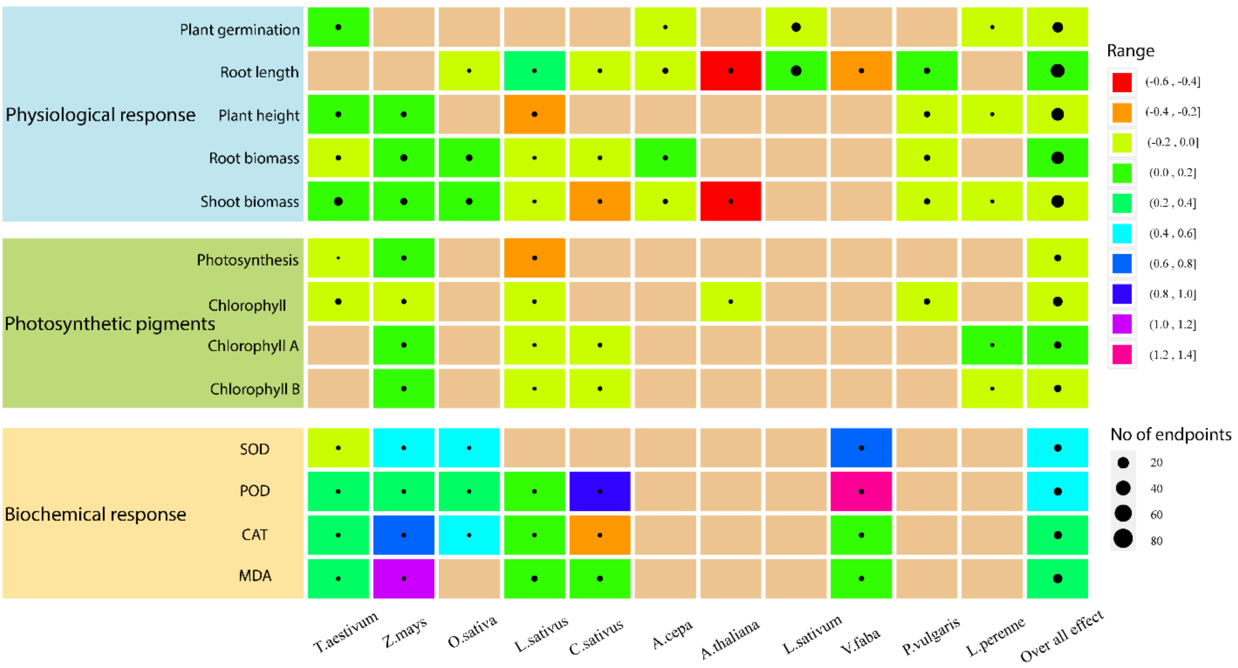


**Figure S4.** Ecotoxicological effects of MPx/NPx on plant species. The circle shows the number of endpoints and the color indicates the effect on the plant.

The increase in antioxidant enzymes indicates that plastic exposure activates defense systems in most species, with generally significant and consistent increases in SOD, POD, CAT, and MDA; the exceptions are SOD in *T. aestivum* (12% reduction) and POD (18% reduction) in *C. sativus*. Antioxidant defense systems protect plant cells and biomolecules from oxidative damage by quenching ROS generated in response to abiotic or biotic stressors.[15] The POD, CAT, and MDA content were increased in *T. aestivum* upon PS-NPx exposure at 10 mg $L^{-1}$ with Cd, whereas the SOD was decreased. The decreased SOD activity may be due to the considerable decrease in Fe, Cu, and Mn content.[16] Whereas, micronutrient are the important components as in SOD isozymes biosynthesis such as Mn-SOD, Fe-SOD, and Cu-SOD, enzymes.[17] Similarly in *Z. mays*, the levels of SOD, CAT, POD, and MDA were increased by exposure to PS-NPx at 1 mg $L^{-1}$.[18] Furthermore, similar response of antioxidant enzymes were observed in *V. faba* and *C. sativus* that were exposed to PS NPx at 100 mg $L^{-1}$ and 50 mg $L^{-1}$ respectively.[3, 10, 19] In summary, MPx/NPx exposure shows negative associations with plant germination, height and shoot biomass, but generally non-significant impacts on root length and biomass. For photosynthesis, all endpoints (photosynthetic rate, chlorophyll b, and total chlorophyll contents) were generally negatively affected, with the exception being chlorophyll a; whose contents increased upon exposure to

MPx/NPx whereas, antioxidant enzymatic activity was significantly increased in different plant species.

(12) Qi, Y.; Yang, X.; Pelaez, A. M.; Huerta Lwanga, E.; Beriot, N.; Gertsen, H.; Garbeva, P.; Geissen, V. Macro- and micro- plastics in soil-plant system: Effects of plastic mulch film

residues on wheat (*Triticum aestivum*) growth. *Science of The Total Environment* **2018**, *645*, 1048-1056. DOI: https://doi.org/10.1016/j.scitotenv.2018.07.229.
(13) Rico, C. M.; Peralta-Videa, J.; Gardea-Torresdey, J. Chemistry, biochemistry of nanoparticles, and their role in antioxidant defense system in plants. In *nanotechnology and plant sciences*, Springer, 2015; pp 1-17.
(14) Wang, F.; Zhang, X.; Zhang, S.; Zhang, S.; Sun, Y. Interactions of microplastics and cadmium on plant growth and arbuscular mycorrhizal fungal communities in an agricultural soil. *Chemosphere* **2020**, *254*, 126791.
(15) Gill, S. S.; Tuteja, N. Reactive oxygen species and antioxidant machinery in abiotic stress tolerance in crop plants. *Plant Physiology and Biochemistry* **2010**, *48* (12), 909-930. DOI: https://doi.org/10.1016/j.plaphy.2010.08.016.
(16) Lian, J.; Wu, J.; Zeb, A.; Zheng, S.; Ma, T.; Peng, F.; Tang, J.; Liu, W. Do polystyrene nanoplastics affect the toxicity of cadmium to wheat (*Triticum aestivum L.*)? *Environmental Pollution* **2020**. DOI: 10.1016/j.envpol.2020.114498.
(17) Gill, S. S.; Tuteja, N. J. P. p.; biochemistry. Reactive oxygen species and antioxidant machinery in abiotic stress tolerance in crop plants. **2010**, *48* (12), 909-930.
(18) Sun, H.; Lei, C.; Xu, J.; Li, R. Foliar uptake and leaf-to-root translocation of nanoplastics with different coating charge in maize plants. *Journal of Hazardous Materials* **2021**, *416*, 125854. DOI: 10.1016/j.jhazmat.2021.125854.
(19) Jiang, X.; Chen, H.; Liao, Y.; Ye, Z.; Li, M.; Klobučar, G. Ecotoxicity and genotoxicity of polystyrene microplastics on higher plant Vicia faba. *Environmental Pollution* **2019**. DOI: 10.1016/j.envpol.2019.04.055.
